# Flux balance analysis and peptide mapping elucidate the impact of bioreactor pH on Chinese Hamster Ovary (CHO) cell metabolism and N-linked glycosylation in the Fab and Fc regions of the produced IgG

**DOI:** 10.1101/2024.08.01.606220

**Authors:** Jayanth Venkatarama Reddy, Sumit Kumar Singh, Thomas Leibiger, Kelvin H Lee, Marianthi Ierapetritou, Eleftherios Terry Papoutsakis

**Author notes:** Correspondence: Marianthi Ierapetritou, ORCID # 0000-0002-1758-9777, Eleftherios Terry Papoutsakis, ORCID # 0000-0002-1077-1277.

## Abstract

Chinese Hamster Ovary (CHO) cells were grown at different bioreactor pH conditions to detail how bioreactor pH affects cell metabolism and site-specific N-linked glycosylation of the produced broadly neutralizing anti-HIV IgG monoclonal antibody VRC01. The data show that pH affects cell growth, glucose/lactate metabolism, IgG production rates, nonessential amino acid metabolism and ammonia accumulation. Parsimonious Flux Balance Analysis (pFBA) and Flux Variability Analysis (FVA) provide insight into the effect of pH on core intracellular reactions at the different pH conditions and culture durations. pFBA revealed the contribution of sources for the production of the toxic metabolite ammonia and provided insights into the switch from ammonia production to consumption. It also documented that culture duration and pH alter the complex bimodal patterns (production/uptake) of several essential and non-essential amino acids. The VRC01 IgG has N-linked glycosylation sites in both the Fc region and the Fab region. Site-specific N-linked glycan analysis using glycopeptide mapping demonstrated that pH significantly affects the glycosylation profiles of the two IgG sites. The Fc region glycans were completely fucosylated but did not contain any sialylation. The Fab region glycans were not completely fucosylated but contained sialylated glycans. Bioreactor pH affected both the fucosylation and sialylation indexes in the Fab region and the galactosylation index of the Fc region. However, fucosylation in the Fc region was unaffected thus demonstrating that the effect of pH on site-specific N-linked glycosylation is complex.

## 1. Introduction

Engineered monoclonal antibodies (mAbs) are developed to target specific antigens and are used as therapeutic molecules for diseases such as cancer, autoimmune diseases, and viral infections (Puthenpurail et al., 2021). As of 2021, about 870 biotherapeutic molecules were in clinical development (Mullard, 2021). Moreover, the patents on many glycoproteins are expiring, leading to an increase in the approval of biosimilars and the rise of the biosimilar market (Gherghescu and Delgado-Charro, 2020). One of the major challenges of biosimilar production is matching the glycosylation patterns of the novel therapeutic (Hajba et al., 2018). Chinese Hamster Ovary (CHO) cells are used to produce 60% of the monoclonal antibodies on the market (Dhara et al., 2018). CHO cell process development involves cell line development, media optimization and bioreactor process optimization. Process intensification involves determining the best bioreactor operating conditions. It is widespread practice in the industry to shift the temperature of the culture to maximize production of the mAbs (Aghamohseni et al., 2017; Bollati-Fogolín et al., 2005; Fox et al., 2004; McHugh et al., 2020). However, literature studies have also used pH shifts to maximize the titer and control product quality (Hogiri et al., 2018; Trummer et al., 2006). Scaling up the production to large scale bioreactors shows that gradients in pH within the bioreactor can significantly affect the metabolism and N-linked glycosylation of the cells (Brunner et al., 2017a; Brunner et al., 2017b). Bioreactor pH is known to affect titer production rates, growth rates and nutrient uptake rates (Borys et al., 1993; Yoon et al., 2005). These differences in uptake rates of nutrients can lead to suboptimal media performance at certain pH values. Studies have shown that culture pH can affect N-linked glycosylation of biologics (Borys et al., 1993). More specifically it has been shown that bioreactor pH can affect the galactosylation of glycoproteins (Aghamohseni et al., 2014; Ivarsson et al., 2014). However, the effect of critical process parameters (pH,

temperature, and DO) on site specific N-linked glycosylation of mAbs has not been reported in the literature. This is important because about 20% of the IgG in the human serum contain glycans in the Fab region and the Fc region (Van De Bovenkamp et al., 2016; Xu et al., 2021). The effect of Fc glycosylation on mAb affinity, half-life, aggregation, and thermal stability have been well studied over the past few decades (Liu, 2015). Recent studies also show that Fab glycans can also influence mAb affinity, activity, half-life and aggregation (Van De Bovenkamp et al., 2016). One study introduced an N-linked glycosylation site in the Fab region of a mAb by modifying the amino acid sequence. It was shown that mAbs with glycans in the Fab region had increased half-life in mice and showed similar efficacy to the original molecule (Ludwig et al., 2022). Another study looked at thermal stability of mAbs with and without Fab glycans and showed that the presence of Fab glycans improves thermal stability of antibodies (van de Bovenkamp et al., 2018). Hence there is the need to control Fab N-linked glycan profiles and treat it with the same importance that we treat the Fc glycan fractions. From a Quality by Design (QbD) perspective, it is important to understand the impact of process parameters on site specific N-linked glycan fractions. Although this study focuses on Fab and Fc glycans, the importance of studying the effect of critical process parameters on site specific N-linked glycosylation can extend to any glycoprotein that has several N-linked glycosylation sites.

To study the effect of bioreactor pH on CHO cell metabolism and site-specific N-linked glycosylation, CHO cells producing a broadly neutralizing anti-HIV antibody (VRC01) were cultivated in bioreactors at three different pH conditions. Measurements of glucose, lactate, amino acids, titers, ammonia, and site-specific N-linked glycan fractions at different timepoints allowed us to study the effect of bioreactor pH and culture duration on CHO cell metabolism and product quality. A survey of the literature showed that there are limited number of studies that utilize stoichiometric models to study the effect of bioreactor critical process parameters (pH and DO) on cellular metabolism (Reddy et al., 2023). In the literature, parsimonious Flux Balance Analysis (pFBA) along with constraining the maintenance energy has been used to study the metabolism of CHO cells (Széliová et al., 2021). Another study has used FBA to study the effect of bioreactor temperature shifts on CHO cell metabolism (Sou et al., 2015). Measurements of glucose, lactate, amino acids, titers, and ammonia have allowed us to use a similar approach to explain the differences in metabolic uptake/production rates observed at the various pH values and provide insights into intracellular metabolism. However, using underdetermined system of reactions to make conclusions on the effect of pH on cell metabolism can be tricky, due to the existence of multiple possible solution (Pan et al., 2017). To overcome this challenge, Flux Variability Analysis (FVA) has been used to study the range of possible fluxes. This study utilized experiments and stoichiometric models to study the differences in cell metabolism at different pH conditions and culture durations. The results presented in this study reveal the drastic differences between glycans observed in the Fab and Fc regions of the antibody. It was found that the changes in overall glycosylation indices as a function of bioreactor pH were significant. Changes in glycosylation indices in one site do not always indicate a similar change in the same glycosylation index on the other site. Hence, showing that the N-linked glycan fractions across the two sites are not uniformly affected by changes in bioreactor pH is an unexpected and novel finding. Improving our knowledge of this can help control the site-specific N-linked glycan fractions and aid with implementing quality by design for glycoproteins with multiple N-linked glycosylation sites.

## 2. Materials and methods

### 2.1 Cell culture

CHO-K1 cell line (Clone A11 from the Vaccine Research Center at the National Institute of Health) stably expressing a broadly neutralizing anti-HIV monoclonal antibody (VRC01) with Fab and Fc glycosylation was used in this study. CHO cells were grown in 125 mL Erlenmeyer flasks at 30 mL culture volume for passage and inoculum preparation. HyClone Actipro media (Cytiva) supplemented with 6 mM L-glutamine was used as the basal media. HyClone Cell Boost 7a supplement and HyClone Cell Boost 7b supplement were used as feed media. CHO cells were grown in 1 L Eppendorf bioflo 120 bioreactors at an initial culture volume of 750 mL. The bioreactor was operated at temperature of 37 °C, dissolved oxygen setpoint of 40 % and agitation set to 90 RPM using a pitched blade impeller. Three different bioreactor pH values (6.75, 7, and 7.25) were studied by running biological triplicates for each condition. The bioreactor pH was controlled by sparging with CO2 and pumping 6% sodium bicarbonate solution. A significant amount of base was required only for the pH 7.25 condition due to higher lactate accumulation. The feed 7b medium has a basic pH to improve solubility of tyrosine. Thus, sodium bicarbonate addition was not necessary for the pH of 7 condition. The dissolved oxygen was controlled by sparging with oxygen and air. Starting day 3 of the culture, feed media was added daily, 22.5 mL of HyClone Cell Boost 7a and 2.25 mL of HyClone Cell Boost 7b were added. Starting day 5, glucose was added to the bioreactor every day to bring the glucose concentration to 9 g/L. Antifoam C was added when foaming was observed. Samples were collected daily for measurements of cell counts, glucose concentration, lactate concentration, ammonia concentration, amino acid concentration and titer. Samples were collected every 2 days starting from day 4 for site specific glycan analysis. Cultures were terminated when viability dropped below 80%. The viability dropped below 80% on day 11 or 12 for the bioreactor runs at pH 7 and pH 7.25. Thus, site specific glycan analysis has been compared only from day 4 to day 10 for these two conditions. The titers on day 4 were very low of bioreactor run at pH 6.75. Thus, site specific glycan analysis has been performed from day 6 to day 14 for this condition. Cell counts and viability were measured by using the trypan blue assay on the DeNovix automatic cell counter.

### 2.2 Measurements of substrate, metabolite, and IgG concentrations

Measurements of glucose and lactate were performed using the YSI 2700 bioanalyser. Ammonia concentrations and osmolarity were measured by using the NOVA bioanalyser. Amino acid concentrations were measured using an Agilent HPLC 1260 instrument. Standards and a HPLC column (AdvanceBio Amino Acid Analysis column) were purchased from Agilent. The amino acids were derivatized with OPA for primary amino acids and FMOC for secondary amino acids as per the protocol provided by Agilent. Derivatization was performed on the autosampler. Antibody titers were measured on the Agilent HPLC 1260 using protein A chromatography using a POROS A HPLC column (Catalog number, 1502226; ThermoFischer). The method was performed in accordance with the manufacturer’s protocol. Mobile phase A consisted of 50 mM phosphate and 150 mM NaCl at pH 7.0. Mobile phase B consisted of 12 mM hydrochloric acid at pH 1.9. The gradient of mobile phases is in supplementary Table S2. A UV detector (280 nm) was used to detect the mAbs. Additional method details are provided in the supplementary material. HPLC grade IgG standard (Catalog number MFCD00163923) was purchased from Sigma-Aldrich.

### 2.3 Site-specific glycan analysis

Starting day 4, samples (8 mL) were taken every 2 days for site-specific glycan analysis. Due to the laborious and expensive nature of performing site specific glycan analysis, it was performed for biological duplicates. Protein A purification was performed on these samples using an AKTA Pure and protein A HiTrap mAb select column (catalog number 29497628) purchased from Cytiva. These samples were further concentrated using 10 kDa centrifugal filters. 1 mg of mAb was extracted by removing the appropriate volume and vacuum evaporation. 1 mg of mAb was not available on day 4 for duplicate bioreactors run at pH 6.75 because of low titer values. Hence, samples starting day 6 were used for pH 6.75 bioreactor condition. Duplicate samples were also not available for bioreactors run at pH 7.00 for day 4 and day 8 of the cultures. Hence single samples were used only for these two days. Duplicate samples were used for all the other conditions and timepoints. Trypsin digestion and Glu-C digestion were performed on the mAb. LC-MS was performed using a Waters BioAccord with ACQUITY Premier system equipped with a Waters BEH C18 column (300Å, 1.7 μm, 2.1 mm ×150 mm, catalog number 186003687). Trypsin and Glu-C (ammonium bicarbonate pH 7.8) were selected as digest reagents in the analysis method. An absolute retention time mass tolerance of 0.1 min, and target identification mass tolerance of 10 PPM (20 PPM for fragment match tolerance) were specified. Identification of at least five primary fragment ions was required for confirmation. The glycan modification library used for data processing is given in supplemental Table S3 and S4. Additional LC-MS method parameters are listed in supplemental Table S5. UNIFI software and the modification library were used to determine the glycan structures at both sites. The glycosylation indices were calculated separately for the Fc region data and the Fab region data based on equations from the literature (Ghaffari et al., 2020). The fucosylation index is calculated by dividing the number of fucosylated glycans (F1) by the sum of the fucosylated glycans (F1) and the afucosylated glycans (F0). This is shown in Eq. 1. Only biantennary glycans were observed on this mAb. Hence, the galactosylation index (Eq. 2) was calculated by dividing the sum of galactose in G2 (Galactose is present in both the branches) and G1 (galactose is present on only one branch) species by the total number of potential galactose fractions that can be present in a biantennary glycan (two galactose molecules per branch). G0 represents glycans without galactose. Similarly, S2 represents sialic acid molecules on both the branches, S1 represents sialic acid present on only one branch and S0 represents the absence of sialic acid. The sialylation index (Eq. 3) is defined as the total molecules of sialic acid present divided by the total possible number of sialic acid residues. Biantennary glycans contain two branches and can contain a sialic acid molecule on each branch.

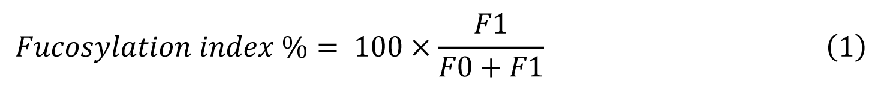

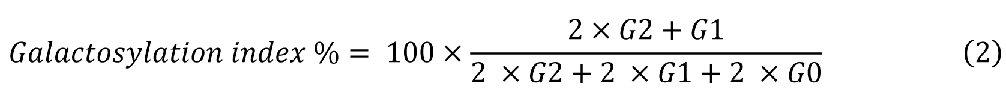

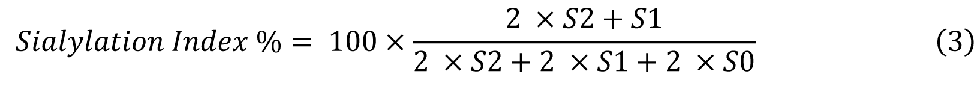

### 2.4 Parsimonious Flux balance analysis (pFBA)

Parsimonious FBA was performed by using a literature network consisting of 144 reactions and 101 metabolites (Jiménez Del Val et al., 2023). This model has been used to study intracellular metabolism at different culture phases. It provides an alternative to genome scale models but contains relevant metabolic information to study CHO cell metabolism. Growth rates, uptake and production rates of amino acids, glucose, cells, titer, lactate, and ammonia have been calculated by plotting the Integral Viable Cell Density (IVCD) vs the cumulative amount consumed or produced (Sou et al., 2015). Glutamine degradation rate has been reported for the Actipro media in the literature (Széliová et al., 2020). The uptake rate of glutamine by the cells is equal to the experimentally observed rate of change in glutamine concentrations divided by the number of cells subtracted by the glutamine degradation rate reported in the literature. Degradation of glutamine results in production of ammonia and pyrrolidonecarboxylic acid (Ozturk and Palsson, 1990). The ammonia production rate was calculated as the experimentally observed change in ammonia concentration divided by the number of cells added to the rate of glutamine degradation. These calculations were made for three different phases of the fed-batch process at the three different pH conditions. The culture durations were split into early growth phase (day 0 to 3), late growth phase (day 4 to 7) and stationary phase (day 8 to 11). The data from biological triplicates were tabulated in Supplementary Table S1. A two-tailed T-test was performed to determine statistical significance between the uptake and secretion rates across the different pH conditions. The following metric was used to depict statistical significance in plots of uptake rates. P values below 0.001 were represented as ***, p values below 0.01 were represented as ** and p values below 0.1 were represented as *. A survey of the literature showed that biomass maximization is the most used objective function to perform flux balance analysis during the growth phase of cultured cells (Jiménez Del Val et al., 2023; Sou et al., 2015; Széliová et al., 2020; Széliová et al., 2021). In fed-batch conditions in which the growth rates, nutrient uptake rates, byproduct secretion rates, and titer production rates are constrained to the experimentally measured value, biomass maximization objective function has been used in the growth phase of the cultures and maximization of specific titer production rates has been used in the stationary phases of culture (Calmels et al., 2019; Sou et al., 2015; Zamorano et al., 2010). Hence, Biomass maximization was used as the objective function for the early growth phase and late growth phase. Maximization of specific titer production rates was used as the objective function for the stationary phase. The lower bounds and upper bounds were determined based on the standard deviation of the experimental measurements for all the measured fluxes. The bounds for all the reversible and irreversible reactions were set based on the information from the literature (Jiménez Del Val et al., 2023). The bounds of fluxes for metabolites that were not measured were set at ±30% of the values reported in the literature (Carinhas et al., 2013). Flux Variability Analysis was also performed to study the range of possible fluxes in each condition with optimality factor set to 0.95 and pFBA factor set to 1.1. pFBA and FVA were performed using cobrapy (Ebrahim et al., 2013). Solutions to underdetermined systems are determined using pFBA. However, there are infinite solutions that a possible for a range of objective functions. Hence, FVA was performed for all the different objective functions used.

## 3. Results and Discussion

### 3.1 Bioreactor pH impacts cell growth rates and peak cell densities

The effect of bioreactor pH on Viable Cell Density (VCD) and growth rates are shown in Figs. 1a & 1b, respectively. The highest peak VCD of 28.8 million cells/mL was achieved at pH 7, followed by 27.1 million cells/mL at pH 7.25 and 18.2 million cells/mL at pH of 6.75. The statistical test performed on the data from biological triplicates at the three pH conditions show that the peak VCD at pH 7 was not significantly higher than the peak VCD at pH 7.25 (p value > 0.1). However, the peak cell densities at pH 7 and pH 7.25 were significantly higher than the value observed at pH 6.75 (p value < 0.001). The fed-batch culture has been divided into three phases. According to the fed-batch feeding protocol in the literature for this cell line, the cells were cultivated in Actipro media (basal media) with 6 mM glutamine, followed by feeding Cell Boost 7a and Cell Boost 7b. Glutamine was not added with the feed media. (Cordova et al., 2023). High growth rates were observed in the first few days (0 to 3; early growth phase) of the culture. This can be attributed to uptake of glutamine. Fig. 1d shows that glutamine was depleted on day 3 of the cultures. Depletion of glutamine leads to the second phase of the culture. For all three pH values, during the second late-growth phase of the culture, nutrients (except glutamine) were not limiting. The carbon source in the late growth phase is glucose but isotope labeling experiments from the literature have also shown that asparagine can drive TCA cycle fluxes in the absence of glutamine (Kirsch et al., 2022). For the pH 7 and pH 7.25 conditions, the VCD started dropping when asparagine was depleted. Asparagine was added in the feed media, but the amount added could not sustain the growth of cells at such large cell densities. Asparagine (Fig. 1g) was depleted before the measurement at the next 24-hour time point. Depletion of both asparagine and glutamine has been shown to reduce growth rates by 90% (Ghaffari et al., 2020). For pH 7 and 7.25, depletion of nutrients led to the third stationary-culture phase. Fig. 1j shows the plot of osmolarity for all the conditions. In the case of pH 6.75, cell death can be attributed to the increase in osmolarity of the culture. Increase in osmolarity from 320 mOsm per kg to 500 mOsm per kg in CHO cell cultures has been shown to reduce growth rates drastically (Alhuthali et al., 2021). The reasons for the increase in osmolarity in the pH 6.75 condition are discussed later. In the early growth phase, the highest growth rate (Fig. 1b) was observed for the pH 7 condition, followed by pH 7.25 and pH 6.75. The growth rates in the late growth phase were similar for all pH conditions. It was surprising to see that growth rates were only affected during the early growth phase. This could be due to the time taken for rewiring of metabolism to adjust for the difference in pH conditions. The peak VCD was observed on different days for all three pH conditions due to differences in growth rates in the early growth phase. Cultures were terminated when cell viability reached 80%. This occurred on different days for the different pH conditions. The differences in viability profiles seen in Fig. 1c can be attributed to nutrient depletion, osmolarity increases and the potential production of growth inhibitory metabolites. The viability dropped below 80% between day 11 and 12 for pH 7 and pH 7.25. For pH 6.75 viability dropped below 80% on day 14. As stated, the feeding media do not contain glutamine. We tested the impact of adding glutamine to the feeding media in shake flask experiments. We found that it resulted in significant increase in ammonia accumulation and drop in cell viability.

**Fig. 1:**
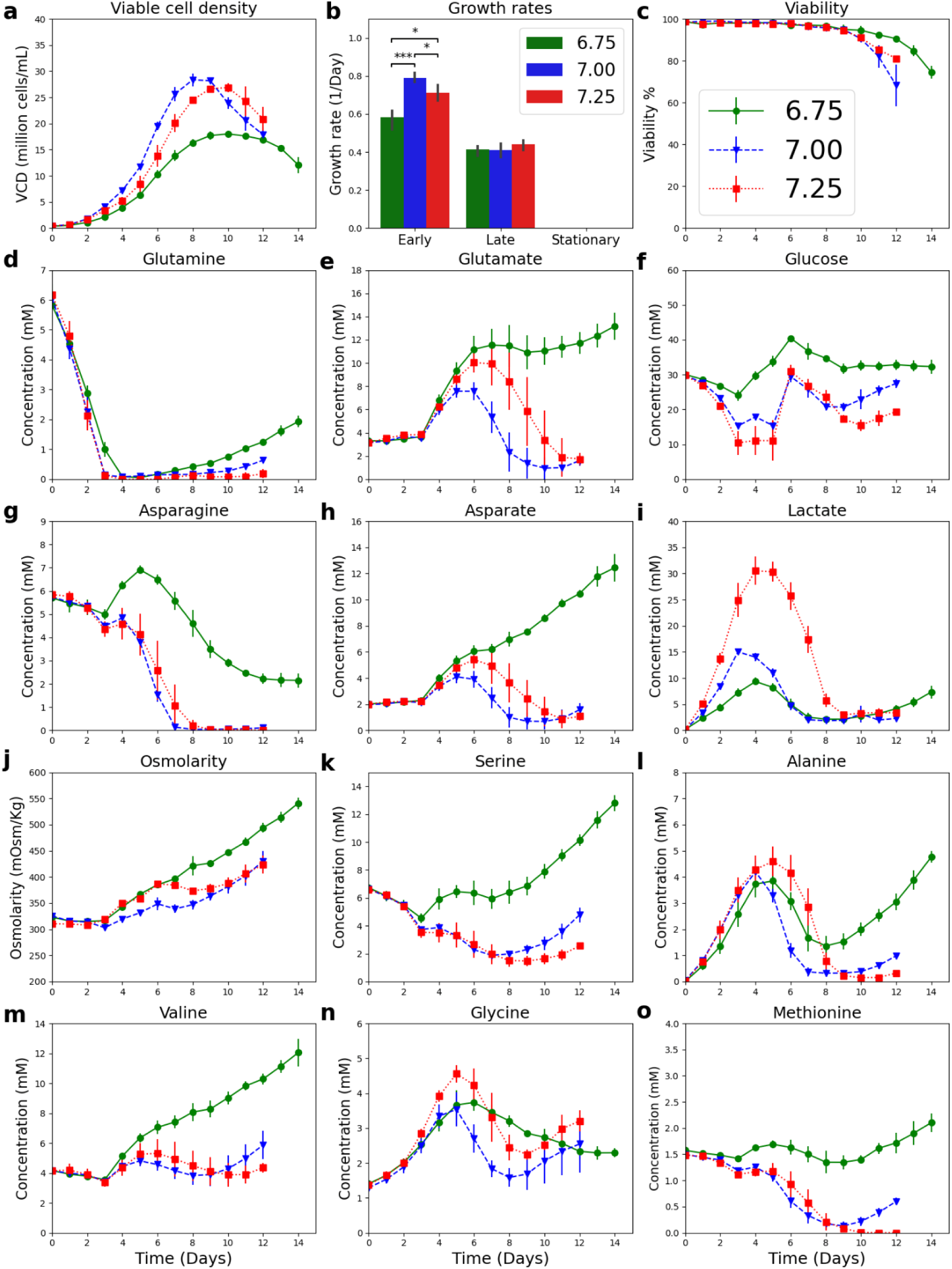
Bioreactor pH affects nutrient depletion and osmolarity thus impacting growth rates, peak viable cell density, time to peak viable cell density, harvest day, cell viability, amino acid concentrations, glucose concentrations and lactate concentrations. Data shown for biological triplicates. a) Viable cell density, b) Growth rate during the early growth phase (day 0 to 3), late growth phase (day 4 to 7) and stationary phase (day 8 to 11), c) Viability, d) Concentration of glutamine, e) Concentration of glutamate, f) Concentration of glucose, g) Concentration of asparagine, h) Concentration of aspartate, i) Concentration of lactate, j) Osmolarity, k) Concentration of serine, l) Concentration of alanine, m) Concentration of valine, n) Concentration of glycine, and o) Concentration of methionine.

### 3.2 Higher bioreactor pH leads to higher rates of glucose uptake and lactate production

Fig. 1f and Fig. S2a show that higher bioreactor pH results in statistically significant increase in cell specific glucose uptake rates. Bioreactor at pH 7.25 showed this trend throughout the culture duration. This trend has been observed by other literature studies (Borys et al., 1993; Yoon et al., 2005). Increased glycolysis flux can be explained as a function of glycolytic enzyme activities and intracellular cytosolic pH. Glycolytic enzymes have been shown to have increased activity at alkaline pH (Alfarouk et al., 2014). Accumulation of intracellular pyruvate coming from glycolysis can lead to production of alanine and lactate as observed in Figs. 1i and 1l. Fig. 1i and supplementary Fig. S2b show that increasing bioreactor pH leads to an increase in cell specific lactate production rates. Switch in lactate metabolism, from production to uptake, has also been reported to be caused by depletion of glutamine. As glutamine enters cell metabolism through the TCA cycle, its depletion can result in reduced TCA cycle fluxes. (Zagari et al., 2013). For pH 7.00 and pH 6.75, the switch in lactate metabolism occurred exactly when glutamine was depleted. Lactate is also known to reduce the pH of the media. Significant accumulation of lactate will require addition of base to control the bioreactor pH. Bioreactor operated at pH of 6.75 did not require addition of base as small amounts of lactate were produced. Bioreactor operated at pH of 7.00 produced a significant amount of lactate. However, addition of the feed media resulted in an increase in pH. Feed 7b has a high pH value as it is required to dissolve tyrosine. No additional base was required for bioreactor operated at pH 7.00. Bioreactor operated at pH of 7.25 required addition of significant amount of base. The consequence of which is an increase in osmolarity of the culture as can be observed in Fig. 1j. If it is desired to operate at high pH conditions, methods to control glycolysis fluxes would result in better culture performance.

### 3.3 Bioreactor pH affects amino acid metabolism and mAb production

Bioreactor pH had a significant effect on uptake and production rates of different amino acids. Glutamine was rapidly consumed by the cells and had the highest uptake rate among all the amino acids. Glutamine is known to be an alternative energy source for rapidly dividing cells (Reitzer et al., 1979). This rapid consumption of glutamine could lead to increased growth rates in the early phase of the culture (Fig. 1). Glutamine supplementation in CHO cells has also been shown to increase the alanine and lactate fluxes (Kirsch et al., 2022). Since glutamine can produce alanine, lactate, aspartate and glutamate, its depletion could have resulted in a switch in metabolism of these metabolites. During the glutamine consumption phase (Fig. 1), lactate, aspartate, glutamate, and alanine were produced. After depletion of glutamine, these metabolites were consumed. An exception to this is the pH 7.25 condition that had increased glycolysis rates that led to a delay in switch of metabolism of lactate and alanine. After glutamine gets depleted, glucose is the main driver of TCA cycle fluxes. Parsimonious Flux balance analysis has been used to understand this metabolism in section 3.4.

Bioreactor pH also significantly affected the metabolism of other non-essential amino acids (Figs. 1 and S2). The metabolism of alanine, glycine and serine is strongly linked to glycolytic fluxes. Glycine, which is not added in the feed media, was produced and consumed by the cells throughout the culture. Glycine is known to be produced from serine in CHO cells (Carinhas et al., 2013). This serine could come from glycolysis or due to serine uptake from the media. Hence, the reasons for switch in glycine metabolism could be attributed to reduction in glucose and serine uptake rates in the different phases of culture. Accumulation of intracellular pyruvate can also result in production of alanine. When comparing bioreactor pH 7.25 to the other pH conditions, unlike lactate, we did not observe a significant increase in alanine production rates early in the culture. The reasons for lack of increased alanine production rates with increased bioreactor pH are explored by using pFBA in section 3.4. We did however notice a delayed switch in alanine metabolism as a function of increased glycolytic fluxes at higher pH and delayed glutamine depletion at pH of 6.75. Serine uptake rates did not vary drastically among the three pH conditions, but they did vary significantly with culture duration. Literature studies have reported serine uptake rates in the range 1.9 fmol/cell/hour to 17 fmol/cell/hour (Ahn and Antoniewicz, 2013; Naik et al., 2023; Yoon et al., 2005). The serine uptake rate here was 31 fmol/cell/hour during the early phase of the culture. High serine fluxes are examined in detail in section 3.4.

Essential amino acid metabolism was strongly correlated to cell growth for all pH conditions (Figs. 1, S1, and S2). Differences in growth rates resulted in different nutritional needs of the cells. Changing the feed amounts could have resulted in improved performance across the different pH conditions. For example, delaying feed addition time and reducing the amount of feed added for the pH 6.75 condition could have led to improved osmolarity of the culture. This could have extended and improved its growth rates. Bioreactors operated at pH 7.25 showed increased amino acid consumption rates towards the end of the culture (Fig. S2), and this led to depletion of methionine (Fig. 1). The data show that optimal feeding patterns for one pH condition are not optimal for another condition. Thus, it is important to recalibrate the fed-batch process and media with changes in bioreactor pH as the uptake rates are affected by pH.

Ammonia is a byproduct of CHO cell metabolism due to amino acid catabolism, and notably that of glutamine and asparagine. Ammonia accumulation inhibits cell growth (Genzel et al., 2005; Yang and Butler, 2000), affects the cell metabolism and the N-linked glycosylation of proteins (Borys et al., 1994; Yang and Butler, 2000). The effect of ammonia on N-linked glycosylation can be attributed, in part at least, to its effect on the pH in the Golgi (Lee et al., 2019). As each N- linked glycosylation enzyme has an optimal pH (Karst et al., 2017). Hence, it is desirable to design the process aiming to minimize ammonia production. Reduction in glutamine and asparagine concentrations in the media has been shown to reduce the ammonia accumulation in the literature (Kirsch et al., 2022; Xu et al., 2014). Cell specific ammonia production rates were very high for the first few days of the culture (Fig. 2b). The second culture phase led to depletion of glutamine, and consumption of glutamate and aspartate. Depletion of glutamine could have resulted in reduced ammonia production. Intracellular production of glutamine and asparagine from glutamate and aspartate requires one ammonia each. This could result in consumption of extracellular ammonia as observed in the experimental data during the second phase of the culture. It is interesting that bioreactor pH significantly affected ammonia production rates in the third phase of the culture. Lower pH leads to an increase in specific ammonia production rates and increase in extracellular ammonia. When ammonia concentrations increase drastically, it has been reported that alanine production is observed (Synoground et al., 2021). Alanine was produced in the early phase of the culture and consumed from day 4 onwards. However, metabolism of alanine went through another switch at pH 6.75. On day 8 of the culture, when ammonia concentration was 10 mM, the cells began producing alanine. In the bioreactor run at pH 7.00, on day 10, when ammonia concentration reached 10 mM there was a switch in alanine metabolism and cells began producing alanine. Ammonia concentrations never exceeded 10 mM at pH 7.25, and thus a shift in alanine metabolism was not observed. The bioreactor run at pH 7.25 had significantly higher amino acid consumption rates but surprisingly lower ammonia production rates. This suggests that the amino acids were used for protein synthesis, thus preventing ammonia production.

**Fig. 2:**
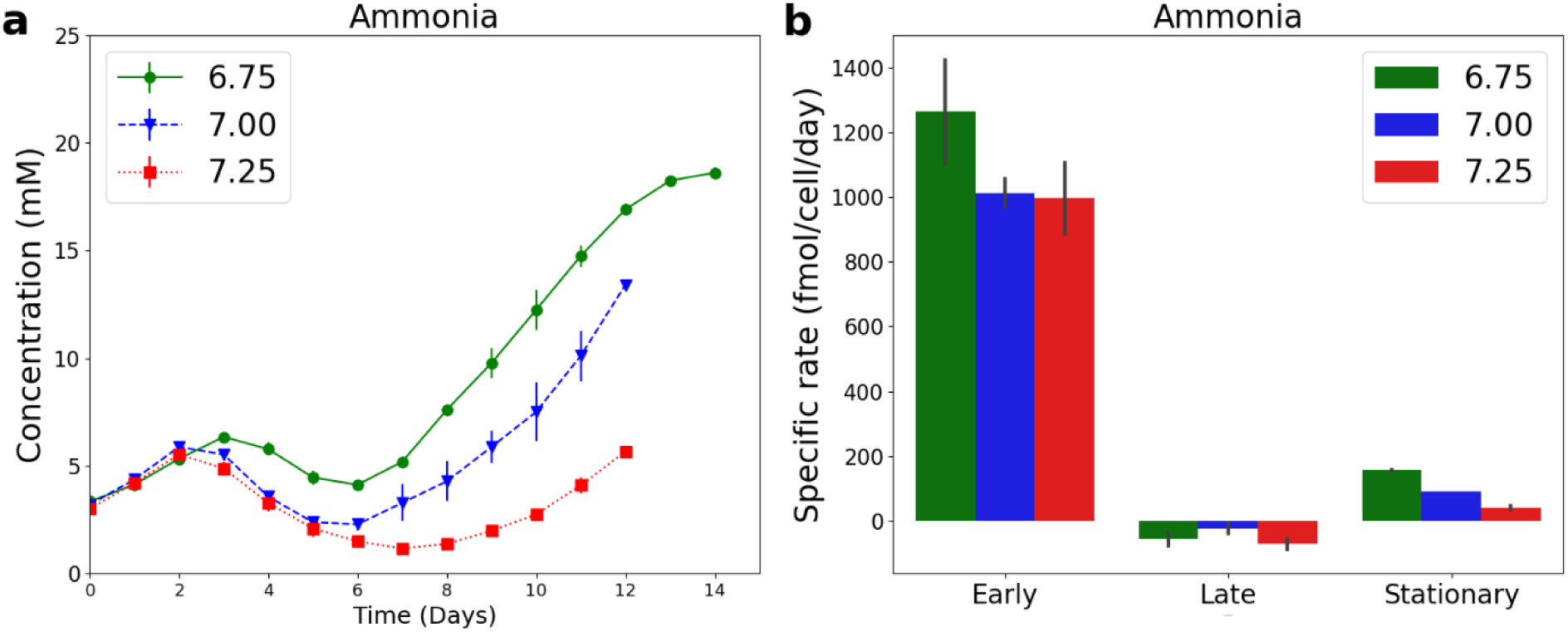
Reduction of bioreactor pH leads to accumulation of ammonia. Ammonia metabolism undergoes multiple shifts in production and consumption rates during fed-batch cultures. Data shown for biological duplicates. a) Concentration of ammonia and b) Cell specific uptake/production rates of ammonia.

Boomi. 3 shows that increase in bioreactor pH led to increased mAb concentrations and increased specific titer production rates. Using transcriptomic and proteomic tools, it was shown that, in the range of 6.9 to 7.1, higher pH leads to upregulation of secretory pathways (Lee et al., 2021). This upregulation can be used to explain the increase in mAb production rates. It can be observed that the best cell specific mAb production rate (qP) is not necessarily observed at the optimal growth rate. Thus pH-shift strategies can be explored for optimizing cell growth and mAb production.

### 3.4 Elucidating the effect of bioreactor pH and culture duration on cellular metabolism using parsimonious Flux Balance Analysis (pFBA)

Stoichiometric models that are constructed from knowledge of metabolic pathways can be used to study the intracellular metabolism of the cell. These models have been used to study the effect of sparging stress, culture duration, temperature and media composition on CHO cell metabolism (Hong et al., 2020; Naderi et al., 2011; Quek et al., 2010; Sou et al., 2015). However, there are limited studies in the literature that use these models to study the effect of bioreactor parameters such as pH or dissolved oxygen on CHO cell metabolism. As demonstrated above (Figs. 1-3), the effect of bioreactor pH on CHO cell metabolism is complicated and can result in switch in metabolism of metabolites such as ammonia, lactate, aspartate, glutamate, glycine, and alanine. To gain more insights into the effect of pH on CHO cell metabolism, a combination of pFBA and FVA were used.

**Fig. 3:**
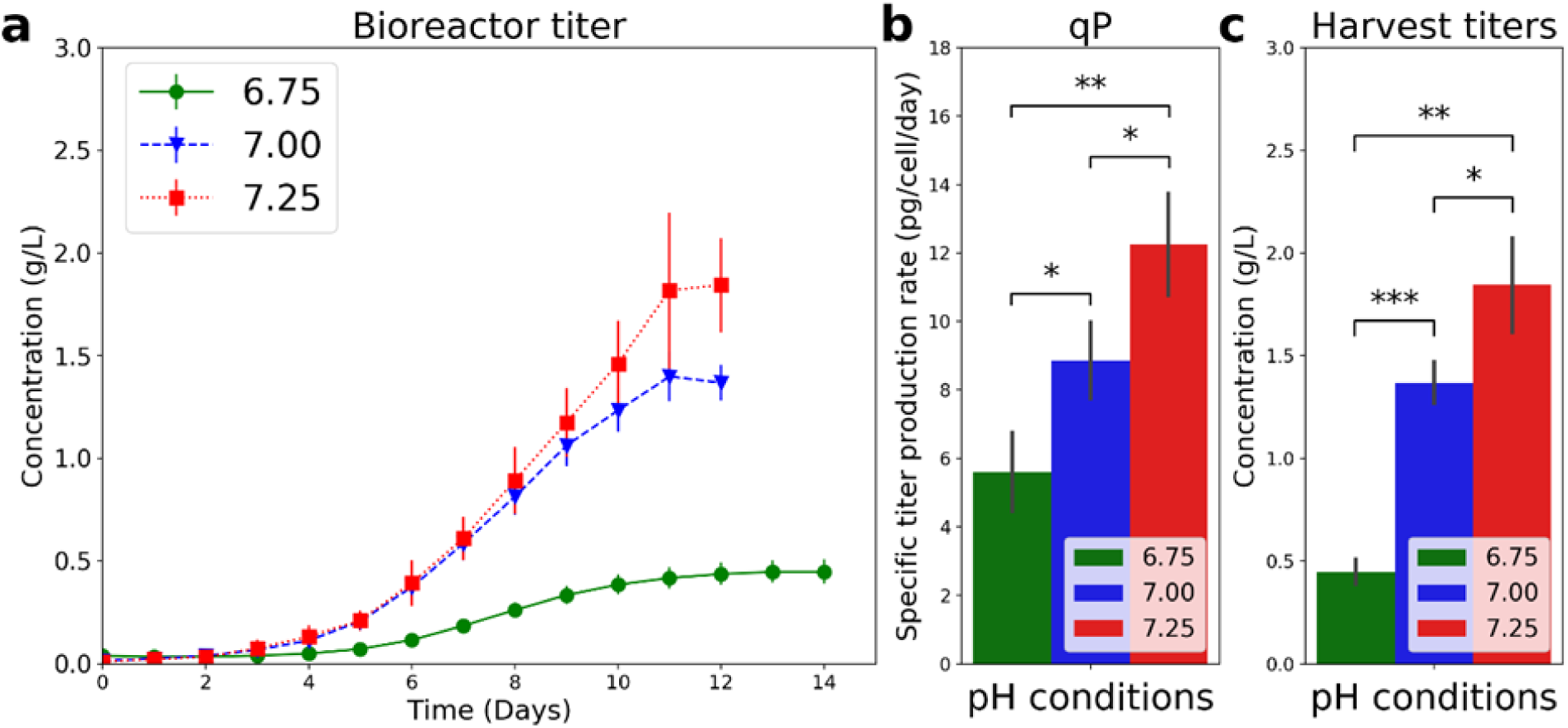
Bioreactor pH affects cell specific antibody production rates. Changes in cell-specific antibody production rates and the viable cell density led to differences in mAb titers. Data shown for biological triplicates. a) Concentration of VRC01 mAb, b) Cell specific titer production rates (qP) and c) mAb titers at harvest.

pFBA analysis (Fig. 4) confirmed that during early culture phases, higher pH increased the flux from glucose to pyruvate. The highest glycolysis fluxes throughout the culture duration were observed at pH 7.25. As expected, increased glycolytic fluxes resulted in increased lactate- production fluxes, but this same trend was not observed for alanine production. This is most likely due to the requirement of glutamate to provide nitrogen to convert pyruvate to alanine and bioreactor pH did not significantly affect glutamine (the main source of glutamate) consumption rates. Glutaminolysis is the conversion of glutamine to glutamate that is incorporated into the TCA cycle via AKG and leaves the TCA cycle at malate to produce pyruvate and may eventually end up as lactate (Young, 2013). This explains why the upper half of the TCA cycle fluxes were much lower than the lower half of the TCA cycle fluxes (Figs. 4a, 4d, 4g and S5). In section 3.2, we noted that the switch from lactate production to lactate uptake coincided with glutamine depletion, indicating that glutamine was converted to lactate. The results from pFBA add further insights into this possibility, indicating that glutamine was responsible for pyruvate production that resulted in lactate production. In addition to effecting lactate metabolism, reduction of the pyruvate production fluxes, and intracellular glutamate concentrations could have also resulted in switch in metabolism of alanine. During the initial phase of the culture, aspartate and glutamate were being produced (Figs. S2g and S2i), the latter from glutamine conversion. Aspartate production can be linked to two sources, consumption of asparagine and production from oxaloacetate produced from glutaminolysis. All subfigures in Fig. 4 show that there is a constant production of aspartate from asparagine, but there is a drastic reduction in production of aspartate from oxaloacetate after depletion of glutamine indicating that the reduction of glutaminolysis led to reduction in intracellular production of aspartate and this led to consumption of aspartate from the media. Thus, the switch in aspartate metabolism can also be linked to depletion of glutamine. During the early phase of the culture, glucose and glutamine were the two major sources of TCA cycle fluxes. Upon depletion of glutamine, the glycolytic flux feeds the TCA cycle fluxes (Figs. 4b, 4c, 4e, 4f, 4h, 4i and S5) and as a result the fluxes in the upper half are similar to the lower half fluxes of the TCA cycle. Analysis of the data of Figs. 1 and S2 in section 3.3, shows that this CHO-cell line exhibits unusually high cell specific rates of serine consumption. pFBA analysis showed that intracellular serine was produced from serine uptake from the media and from reactions involving glycolytic intermediates and glutamate (Fig. 4 and Fig. S3). pFBA analysis also showed that the intracellular serine is converted to pyruvate, ammonia and glycine (Fig. 4).

**Fig. 4:**
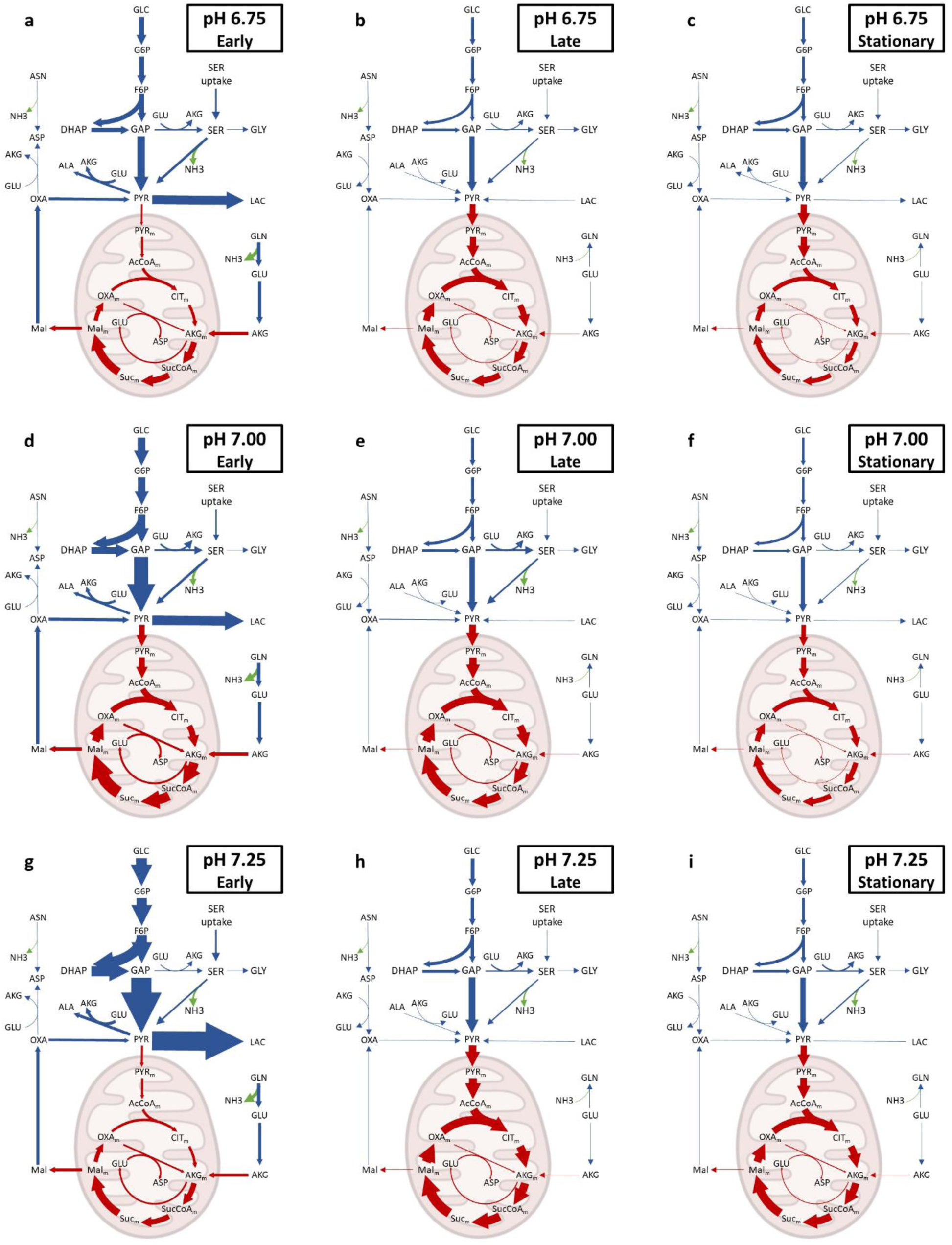
Flux balance analysis reveals major sources of ammonia production and differences in intracellular fluxes at various culture phases for different bioreactor pH conditions.

The switch in ammonia metabolism (Fig. 2) is complicated as it results from the metabolism of several amino acids. Ammonia production reactions (Figs. 4 and S4) indicate that in the early phase of the culture, most of the ammonia is produced from the glutamine conversion to glutamate. Surprisingly the second largest source of ammonia production is not from asparagine but from serine producing pyruvate. pFBA shows that the reaction converting serine to pyruvate led to the largest ammonia production flux in the second and third phase of the cultures. Hence, reduction of serine concentrations in the media could have resulted in better culture performances for product quality and increased titers.

### 3.5 Bioreactor pH impacts Fab and Fc glycosylation

Glycopeptide mapping revealed major differences in Fab and Fc glycans on the IgG1 VRC01 antibody (Fig. 5a). It is not surprising that the Fc region has very high fucosylation and the most abundant glycans were the G0F and G1F. Low amount of high mannose, G0, G2F, and G0F+GlcNAc glycans were also detected in the Fc region. The lack of G2F and sialylation indicates that the galactosyltransferases and sialyltransferase enzymes cannot easily access the glycosylation site in the Fc region. A survey of the glycosylation of over 150 mAbs approved by FDA by May 2023 has been published in the literature. Out of the 150 approved mAbs, it has been reported that 57 IgG1 molecules with glycans only in the Fc region had very similar N-linked glycosylation profiles. G0F glycans were the most abundant followed by G1F and G2F glycans. All 57 IgG1s had very high fucosylation (Luo and Zhang, 2024).

**Fig. 5:**
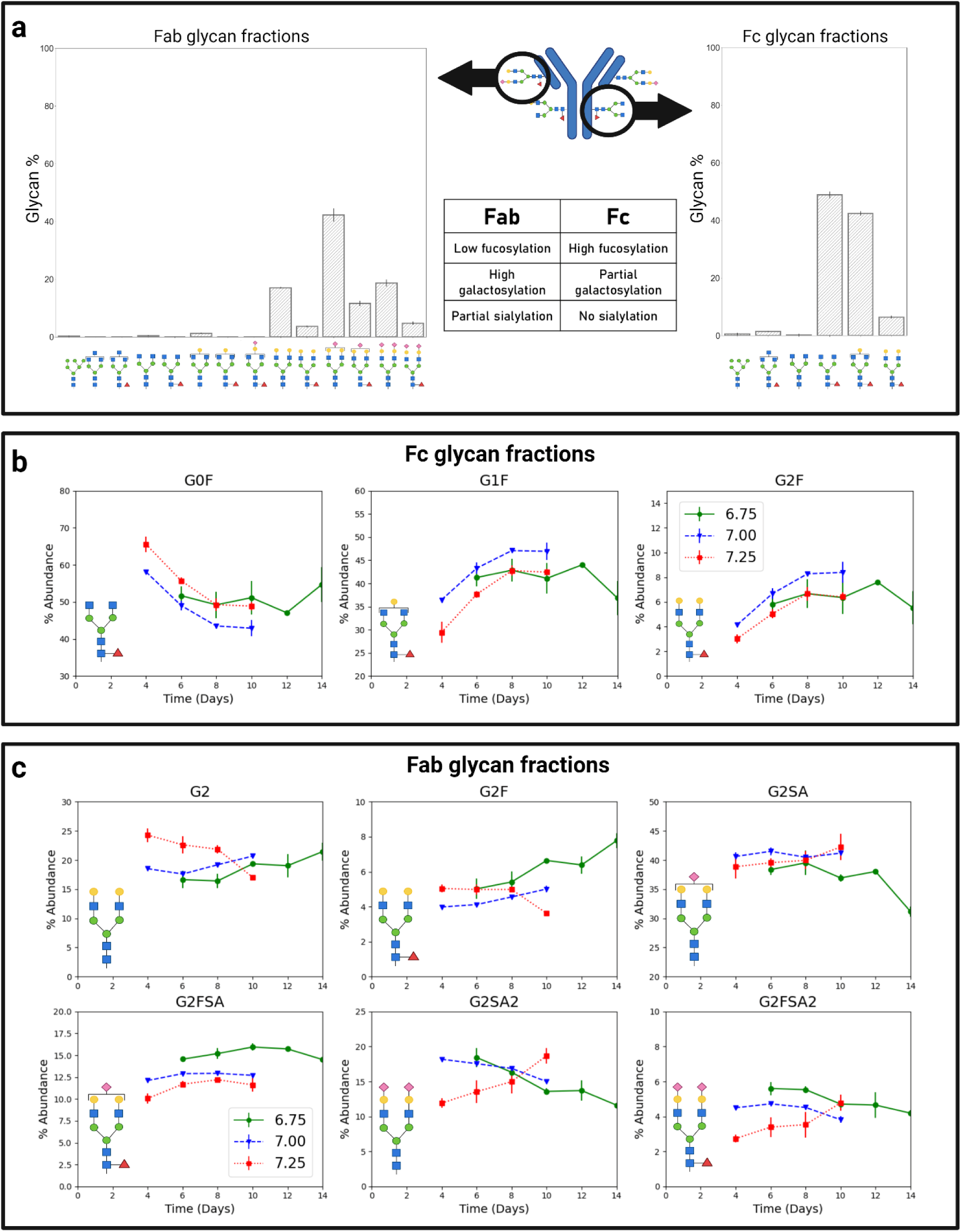
a) Glycopeptide mapping revealed significant differences in the glycans on the Fab and Fc mAb regions. Fab region glycans displayed large diversity, low fucosylation, high galactosylation and partial sialylation. Fc glycans displayed little diversity, high fucosylation, partial galactosylation and no sialylation. Data shown for biological duplicates. Data shown for harvest glycans from bioreactor operated at pH 7.25. b) Glycan fractions in the Fc region were tracked over time for each pH condition. c) Glycan fractions in the Fab region were tracked over time for each pH condition. Glycan fractions of glycans with low abundance are provided in the Supplementary data.

More complex glycoforms were observed in the Fab region while compared to the Fc region (Fig. 5a). The most abundant glycans were G2, G2F, G2S1. G2FS1, G2S2, and G2FS2. Glycans with high degree of sialylation and galactosylation were observed in the Fab region. Hence, indicating increased accessibility of galactosyltransferase and sialyltransferase enzymes to the glycosylation site in the Fab region. The majority of glycans in the Fab region were afucosylated. This implies that for the mAb used in this study, fucosyltransferase enzyme had poor accessibility to this glycosylation site. 20% of IgGs in human serum have glycans in the Fab and Fc region. In the literature site-specific glycan analysis has been performed on IgGs in human serum to identify the differences in glycan fraction across the two sites. It has been reported that Fab glycans contained higher percentages of ternary glycan structures, galactosylation and sialylation but contained lower fucosylation while compared to the glycans present in the Fc region. (Van De Bovenkamp et al., 2016). Site specific N-linked glycosylation on IgG acquired from intravenous immunoglobulin from healthy donors has shown that the fucosylation in Fab region was close to 80% and greater than 95% in the Fc region (Anumula, 2012). The results in our study agree with the literature that the Fab region leads to increased galactosylation and sialylation. However, there were a few significant differences in findings in the literature to IgG from human serum vs. the VRC01. The fucosylation of Fab glycans for the VRC01 was less than 30%. This value is smaller than that reported in the literature (80%) for IgG acquired from intravenous immunoglobulin of healthy donors (Anumula, 2012). The high mannose type and ternary glycan structures were also present in small quantities, indicating good accessibility of mannosidases and poor accessibility of *N*- Acetylglucosaminyltransferase-IV (GnT4) and *N*-Acetylglucosaminyltransferase-V (GnT5) enzymes to the Fab N-linked glycosylation site.

Experimentally determination of the effect of bioreactor pH and culture duration on Fab and Fc glycosylation fractions was an important goal of our study (Figs. 5b and 5c). Bioreactor pH significantly impacted G0F, G1F, and G2F glycans in the Fc region (Fig. 5b). The highest levels of G1F and G2F glycans were observed at pH 7.00. Increase in G1F and G2F glycans naturally result in lower G0F glycoforms. Increasing bioreactor pH led to reduction in G2FSA glycan fractions throughout the culture duration (Fig. 5c). For the pH 7.25 condition, increase in culture time led to reduction in G2 and G2F fractions but increase in G2SA, G2FSA2, and G2SA2 fractions. The opposite trend was observed for these glycan fractions in the pH 6.75 and pH 7.00 conditions. To sum up, bioreactor pH can significantly affect the glycan fractions in the Fab and Fc region. The other glycan fractions present in lesser abundance are shown in Figs. S6 and S7.

Glycosylation indices (as described in Eqs. 1-3) provide a succinct summary of the impact of culture conditions and duration on protein glycosylation. Bioreactor pH had a significant effect on the fucosylation in the Fab region but did not affect the fucosylation in the Fc region (Fig. 6). Fucosylation in the Fc region was nearly 100% throughout the culture duration for all the different pH conditions. Culture duration did not have a significant effect on the fucosylation index of the Fab region. However, an increase in bioreactor pH led to reduction in the fucosylation index. The galactosylation index of the Fc region was the highest at pH 7.00 throughout the culture duration.

**Fig. 6:**
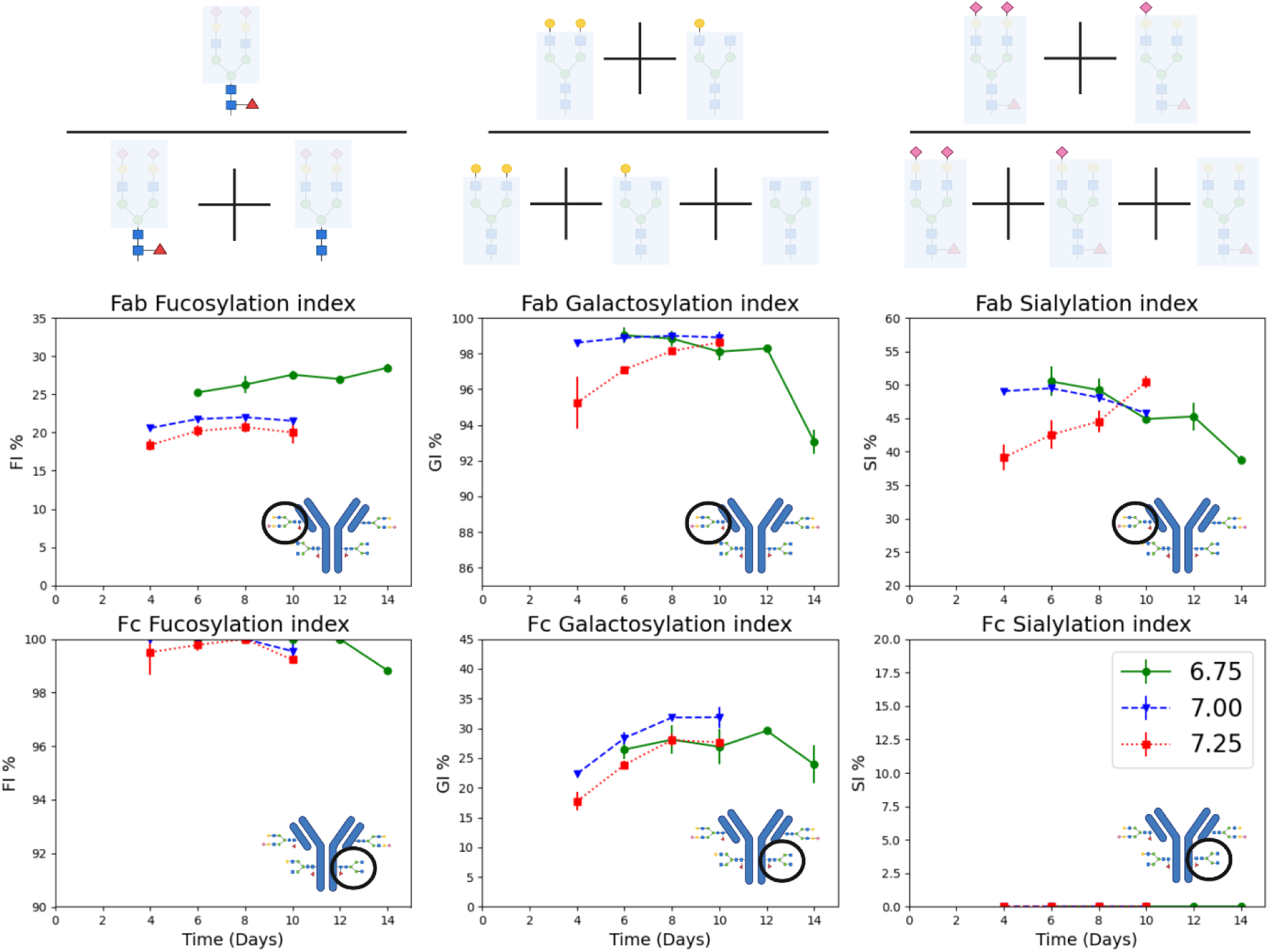
Fucosylation increased with decreasing bioreactor pH in the Fab region but did not change in the Fc region. Bioreactor pH had minimal effects on galactosylation in the Fab region but affected the galactosylation in the Fc region. Sialylation was absent in the Fc region but varied with bioreactor pH and culture duration in the Fab region. Data are presented using glycosylation indices described in Equations (1), (2) and (3). Plots of the effect of bioreactor pH on all the individual glycan fractions are provided in the Supplementary data. Data shown for biological duplicates.

The galactosylation index in the Fc region increased from day 4 to day 8 for pH 7.00 and 7.25. The galactosylation index of the Fab region was very high throughout the culture duration. However, at pH 6.75 the galactosylation index in the Fab region dropped at the end of the culture (Fig. 6). The sialylation index in the Fc region was zero as sialylation was not observed in the Fc region. The effect of bioreactor pH on Fab sialylation was non intuitive. For bioreactors run at pH 7.00 and pH 6.75, the sialylation index of the Fab region dropped as culture duration progressed.

However, the opposite trend was observed for bioreactors operated at pH 7.25. On day 4, there was a drastic reduction in sialylation in the pH 7.25 condition while compared to the other bioreactor runs. The Fab sialylation index in this condition increased as culture progressed. Data in the literature suggest that sialyltransferase activity is most sensitive to changes in Golgi pH (Karst et al., 2017) and that the concentrations of ammonia significantly affect Golgi pH (Lee et al., 2019). High lactate concentrations have also been shown to impact intracellular pH (Lee et al., 2019). Figs. 1 and 2 show that bioreactor pH had a significant effect of lactate and ammonia concentrations.

## 4. Conclusions

Our results indicate that bioreactor pH influences growth rates, glucose uptake rates, lactate production rates, amino acid uptake rates, ammonia secretion rates, and titer production rates. Indicating that the bioreactor pH should not be neglected while designing a fed-batch process and the media (basal and feed) will not be optimal for all the different bioreactor pH conditions. Differences in growth rates and nutrient uptake rates will lead to accumulation or depletion of nutrients at certain pH conditions. Taking this into account can improve product titers. pFBA showed that depletion of glutamine coincides with a metabolic shift from lactate production to lactate consumption. This analysis also showed us that bioreactor pH significantly affects glycolysis and intracellular pyruvate production rates but did not affect alanine production rates in the early culture phase. The effect of glutamine uptake rates on the shift in metabolism of aspartate and glutamate was also dissected by pFBA. We have shown that the switch in metabolism of aspartate is linked to glutamine uptake rates. Bioreactor pH and culture duration significantly affected ammonia metabolism. This CHO cell line also exhibited very high serine uptake rates. Analysis of our data suggests that serine produces pyruvate and leads to ammonia production.

This is the first study to report the effect of a critical process parameter (pH), on site specific N-linked glycosylation of a therapeutic glycoprotein. Our data demonstrated the diversity of glycans present in the Fab and Fc region of the mAb. Fc glycans contain high fucosylation, moderate galactosylation and no sialylation. Fab glycans contained many more species but contained moderate fucosylation, high galactosylation and moderate sialylation. Increased pH led to reduction in fucosylation in the Fab region but did not affect fucosylation in the Fc region. In the Fc region, the highest galactosylation was observed at pH 7. However, bioreactor pH did not seem to drastically affect Fab galactosylation. Sialylation in the Fab region seemed to be affected by metabolites that can impact the intracellular pH. Certain glycosylation attributes can be affected in one site but not the other site. Understanding the effect of bioreactor pH on CHO cell metabolism and site-specific N-linked glycosylation can improve bioprocess performance and implement quality by design.

## Acknowledgements

This research was funded by US FOOD and DRUG ADMINISTRATION, grant number DHHS- FDA-R01FD006588, grant number U01FD007695A. This research was also supported in part by the National Institute of Standards and Technology (70NANB17H002).

## Abbreviations

CHO: Chinese Hamster Ovary
mAb: monoclonal Antibody
pFBA: parsimonious Flux Balance 2 Analysis
FVA: Flux Variability Analysis
QbD: Quality by Design
DO: Dissolved Oxygen
FI: Fucosylation Index
GI: Galactosylation Index
SI: Sialylation Index
VCD: Viable Cell Density
TCA: tricarboxylic acid
AKG: α-Ketoglutarate
MFA: Metabolic Flux Analysis
IVCD: Integral Viable Cell Density

