## Supplementary tables and figures for "Flux balance analysis and peptide mapping elucidate the impact of bioreactor pH on Chinese Hamster Ovary (CHO) cell metabolism and N-linked glycosylation in the Fab and Fc regions of the produced IgG"

Table S1: This table contains data on uptake and secretion rates calculated from the transient metabolic profile data shown in figure 1. This data has been used to perform flux balance analysis. The units of the uptake and secretion rates are in fmol/cell/day. The units for the growth rate or biomass production rate is in 1/day.

| Metabolite | pH 6.75 | | | pH 7 | | | pH 7.25 | | |
|---|---|---|---|---|---|---|---|---|---|
| | Day 0 to 3 | Day 4 to 7 | Day 8 to 11 | Day 0 to 3 | Day 4 to 7 | Day 8 to 11 | Day 0 to 3 | Day 4 to 7 | Day 8 to 11 |
| Biomass | 0.58 ± 0.045 | 0.41 ± 0.03 | 0.0 ± 0.0 | 0.79 ± 0.019 | 0.41 ± 0.024 | 0.0 ± 0.0 | 0.71 ± 0.033 | 0.44 ± 0.022 | 0.0 ± 0.0 |
| GLC | -2327 ± 586 | -1301 ± 47 | -1037 ± 82 | -3420 ± 150 | -1364 ± 72 | -1105 ± 99 | -4941 ± 300 | -1669 ± 88 | -1306 ± 25 |
| LAC | 3096 ± 601 | -229 ± 69 | 24.11 ± 9 | 4167 ± 126 | -254 ± 21 | 1.21 ± 7 | 7255 ± 982 | -199 ± 70 | -24 ± 24 |
| ALA | 1002 ± 111 | -112 ± 22 | -1.7 ± 4.2 | 955 ± 85 | -117 ± 12 | -13 ± 0.7 | 1051 ± 194 | -56.9 ± 12 | -25 ± 2.3 |
| GLY | 456 ± 27 | 44 ± 8 | -7 ± 8 | 318 ± 44 | -15 ± 2 | 13 ± 6 | 404 ± 59 | 22 ± 21 | 11 ± 10 |
| SER | -769 ± 23 | -270 ± 29 | -62 ± 15 | -709 ± 99 | -207 ± 10 | -58 ± 3 | -740 ± 56 | -266 ± 31 | -76 ± 2 |
| GLN | -1776 ± 244 | 5.68 ± 6 | 13 ± 1 | -1773 ± 145 | 2 ± 1 | 4 ± 1 | -1837 ± 179 | 2 ± 2 | 0 ± 1 |
| GLU | 162 ± 16 | -56 ± 39 | -122 ± 19 | 167 ± 18 | -143 ± 18 | -109 ± 12 | 344 ± 77 | -68 ± 24 | -169 ± 11 |
| ASN | -202 ± 36 | -215 ± 9 | -133 ± 15 | -279 ± 54 | -227 ± 10 | -65 ± 2 | -315 ± 20 | -268 ± 28 | -67 ± 2 |
| ASP | 129 ± 11 | -63 ± 6 | -15 ± 17 | 68 ± 10 | -98 ± 11 | -55 ± 2 | 129 ± 13 | -58 ± 20 | -87 ± 7 |
| MET | -45 ± 24 | -41 ± 2 | -11 ± 4 | -73 ± 14 | -43 ± 3 | -9 ±1 | -80 ± 36 | -45 ± 1 | -15 ± 2 |
| TYR | -51 ± 3 | -44 ± 8 | -13 ± 5 | -79 ± 17 | -48 ± 5 | -13 ± 1 | -65 ± 12 | -50 ± 4 | -25 ± 2 |
| PHE | -74 ± 19 | -52 ± 5 | -15 ± 9 | -88 ± 16 | -59 ± 3 | -20 ± 1 | -69 ± 23 | -65 ± 2 | -36 ± 2 |
| TRP | -86 ± 33 | -18 ± 1 | -7 ± 2 | -63 ± 24 | -23 ± 3 | -7 ± 1 | -85 ± 34 | -25 ± 1 | -11 ± 1 |
| VAL | -145 ± 12 | -86 ± 10 | -44 ± 10 | -178 ± 17 | -110 ± 11 | -40 ± 3 | -151 ± 27 | -115 ± 8 | -63 ± 3 |
| LEU | -237 ± 42 | -106 ± 12 | -38 ± 12 | -274 ± 90 | -129 ± 11 | -43 ± 1 | -243 ± 36 | -140 ± 4 | -66 ± 5 |
| ILE | -155 ± 17 | -76 ± 6 | -26 ± 10 | -144 ± 32 | -65 ± 23 | -27 ± 4 | -150 ± 16 | -93 ± 3 | -44 ± 2 |
| ARG | -118 ± 20 | -54 ± 5 | -13 ± 7 | -136 ± 6 | -71 ± 5 | -13 ± 1 | -132 ± 3 | -71 ± 2 | -26 ± 5 |
| HIS | -52 ± 18 | -29 ± 3 | -9 ± 2 | -81 ± 29 | -33 ± 3 | -9 ± 1 | -56 ± 12 | -34 ± 2 | -14 ± 2 |
| THR | -106 ± 49 | -57 ± 6 | -19 ± 11 | -150 ± 52 | -81 ± 7 | -22 ± 1 | -112 ± 25 | -87 ± 2 | -41 ± 4 |
| LYS | -139 ± 9 | -73 ± 7 | -17 ± 10 | -201 ± 47 | -94 ± 7 | -20 ± 1 | -176 ± 10 | -95 ± 3 | -36 ± 5 |
| PRO | -254 ± 88 | -145 ± 21 | -47 ± 9 | -288 ± 98 | -142 ± 17 | -39 ± 4 | -201 ± 96 | -144 ± 15 | -52 ± 8 |
| AMM | 1265 ± 159 | -55 ± 21 | 158 ± 2 | 1013 ± 42 | -22 ± 15 | 91 ± 1 | 997 ± 110 | -71 ± 15 | 40 ± 6 |
| IgG | 0.04 ± 0.005 | | | 0.06 ± 0.006 | | | 0.08 ± 0.008 | | |

Table S2: Quantification of mAb titers using protein A chromatography. Flow rate 3 mL/min. Injection volume 20 μL.

| Time (min) | Gradient (%B) |
|---|---|
| 0 | 0 |
| 0.5 | 0 |
| 0.51 | 100 |
| 1.5 | 100 |
| 1.51 | 0 |
| 3.0 | 0 |

Table S3: Peptide mapping amino acid backbone

| Chain | N-linked glycan site | Peptide sequence |
|---|---|---|
| Light chain | N70 | WGPDYNLTISNLE (65 – 77) |
| Heavy chain | N301 | EEQYNSTYR (297 – 305) |

Table S4: Glycan modification library used in UNIFI for site-specific glycan analysis. Delta mass displayed to the nearest $10^{-4}$ Da.

| Order | Modified name | Delta mass |
|---|---|---|
| 1 | Glycosylation G0F N | 1444.5339 |
| 2 | Glycosylation G1F N | 1605.5867 |
| 3 | Glycosylation G0F-GlcNAc N | 1241.4545 |
| 4 | Glycosylation G1F+SA N | 1897.6281 |
| 5 | Glycosylation G2F+2SA N | 2350.8304 |
| 6 | Glycosylation Man5 N | 1216.4229 |
| 7 | Glycosylation G2F+SA N | 2059.7349 |
| 8 | Glycosylation G2F N | 1768.6395 |
| 9 | Glycosylation G2+2SA | 2204.7724 |
| 10 | Glycosylation G2+SA | 1913.6770 |
| 11 | Glycosylation G2 N | 1622.5816 |
| 12 | Glycosylation G1 N | 1460.5288 |
| 13 | Glycosylation G0 N | 1298.4760 |
| 14 | Glycosylation G0-GlcNAc N | 1095.3966 |
| 15 | Glycosylation G1F-GlcNAc N | 1403.5073 |
| 16 | Glycosylation Man6 N | 1378.4757 |
| 17 | Glycosylation Man7 N | 1540.5285 |
| 18 | Glycosylation Man 8 N | 1702.5813 |
| 19 | Glycosylation Man 9 N | 1864.6342 |
| 20 | Glycosylation Man3-GlcNAc1 | 1095.3966 |
| 21 | Glycosylation Man3-GlcNAc1-Fuc | 1241.4545 |
| 22 | Glycosylation Man3-GlcNAc1-Gal1 | 1257.4494 |
| 23 | Glycosylation Man3-GlcNAc-1-Gal1-Fuc | 1403.5073 |
| 24 | Glycosylation Man3-GlcNAc1-Gal-SA1 | 1548.5448 |
| 25 | Glycosylation Man3-GlcNAc1-Gal-SA1-Fuc | 1694.6027 |
| 26 | Glycosylation Man4-GlcNAc1 | 1257.4494 |
| 27 | Glycosylation Man4-GlcNAc1-Fuc | 1403.5073 |
| 28 | Glycosylation Man4-GlcNAc1-Gal1 | 1419.5022 |
| 29 | Glycosylation Man4-GlcNAc1-Gal1-Fuc | 1565.5601 |
| 30 | Glycosylation Man4-GlcNAc1-Gal1-SA1 | 1710.5977 |
| 31 | Glycosylation Man4-GlcNAc1-Gal1-SA1-Fuc | 1856.6556 |
| 32 | Glycosylation Man5-GlcNAc1 | 1419.5022 |
| 33 | Glycosylation Man5-GlcNAc1-Fuc | 1565.5601 |
| 34 | Glycosylation Man5-GlcNAc1-Gal1 | 1581.5551 |
| 35 | Glycosylation Man5-GlcNAc1-Gal1-Fuc | 1727.6130 |
| 36 | Glycosylation Man5-GlcNAc1-Gal1-SA1 | 1872.6505 |

Table S5: Waters BioAccord LC-MS method parameters used for site-specific glycan analysis.

| | |
|---|---|
| Mobile phase A | Water + 0.1% formic acid |
| Mobile phase B | Acetonitrile + 0.1% formic acid |
| Gradient | **0 to 2 min**, 99.0% A<br>**2 to 65 min**, 99 to 60% A<br>**65 to 68 min**, 60 to 30% A<br>**68 to 70 min**, 30% A<br>**70 to 71 min**, 30 – 99% A<br>**71 to 72 min,** 99% A<br>**72 to 80 min**, 99 to 60% A<br>**80 to 82 min**, 60 to 20% A<br>**82 to 86 min**, 20% A<br>**86 to 87 min**, 20 to 99% A |
| Flow rate (mL/min) | 0.250 |
| TUV wavelength (nm) | 214 |
| RDa mode | Full scan with fragmentation |
| Mass range | Low (50 – 2000 m/z) |
| Scan rate (Hz) | 1 |
| Capillary voltage (kV) | 1.20 |
| Desolvation temperature (°C) | 350 |
| Lockmass | ACQUITY RDa waters connect Lockmass Kit (Waters part number 186009298) |
| Lockmass correction mode | Dynamic |

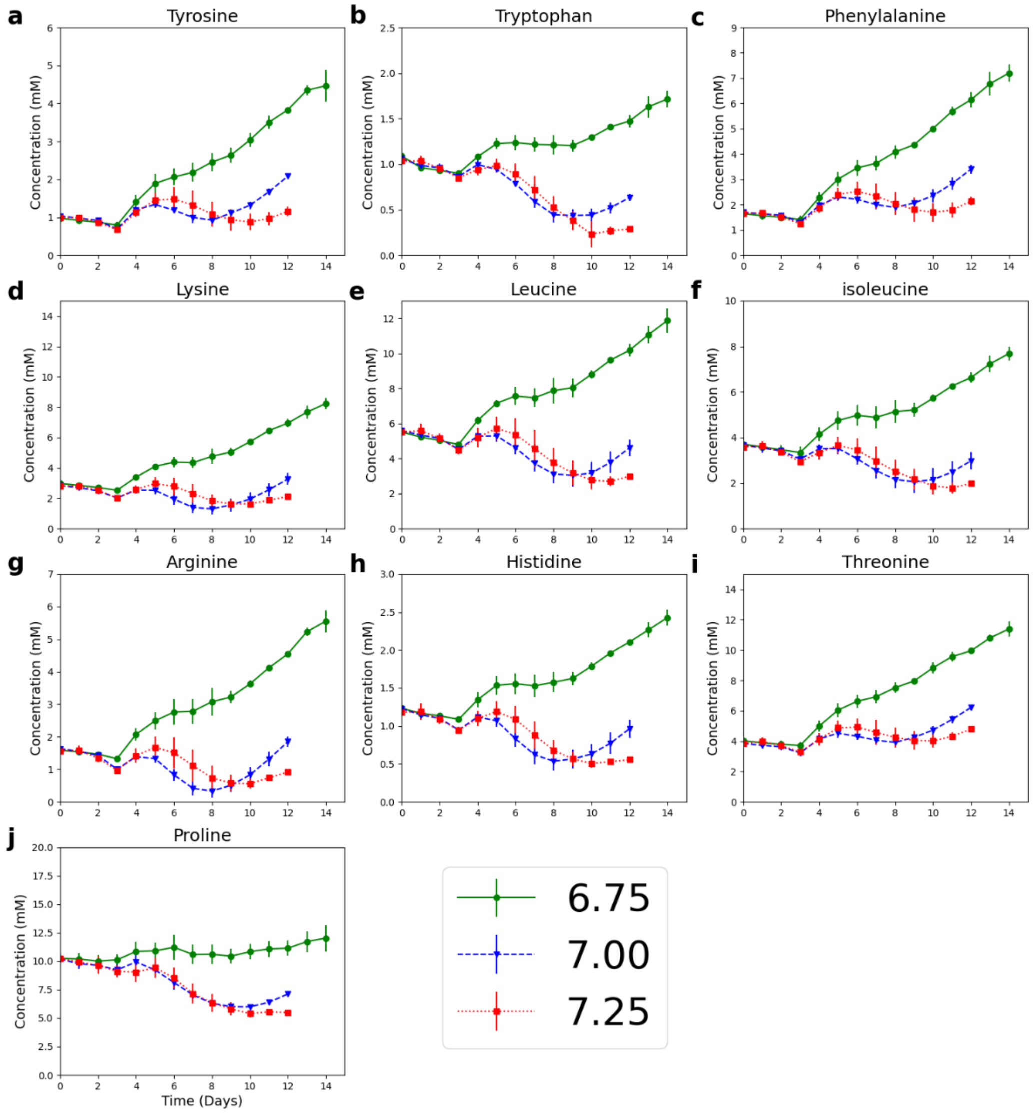


Fig. S1: Effect of bioreactor pH and culture duration on amino acid concentrations.

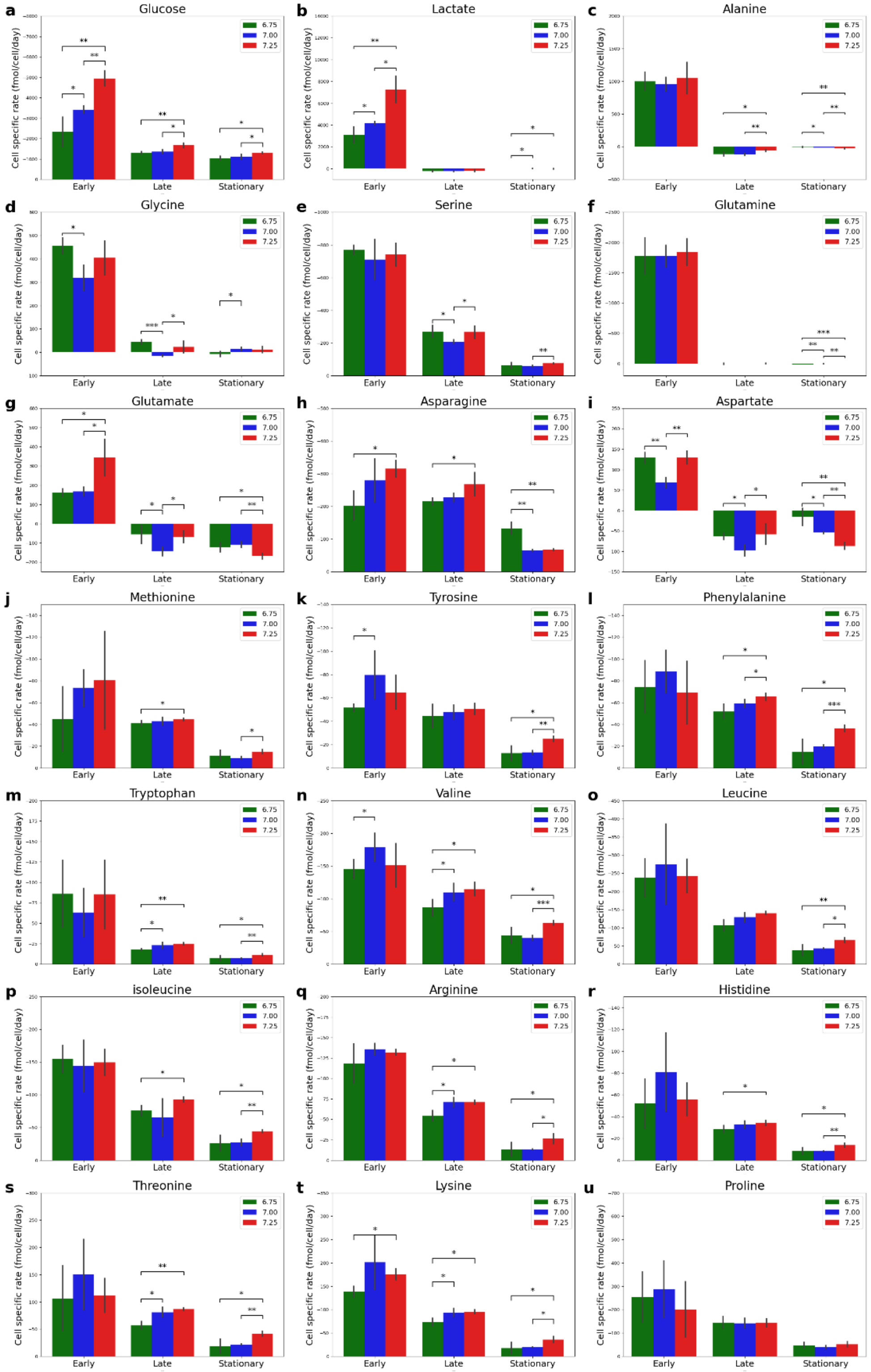


Fig. S2: Effect of bioreactor pH and culture duration on metabolic uptake and production rates.

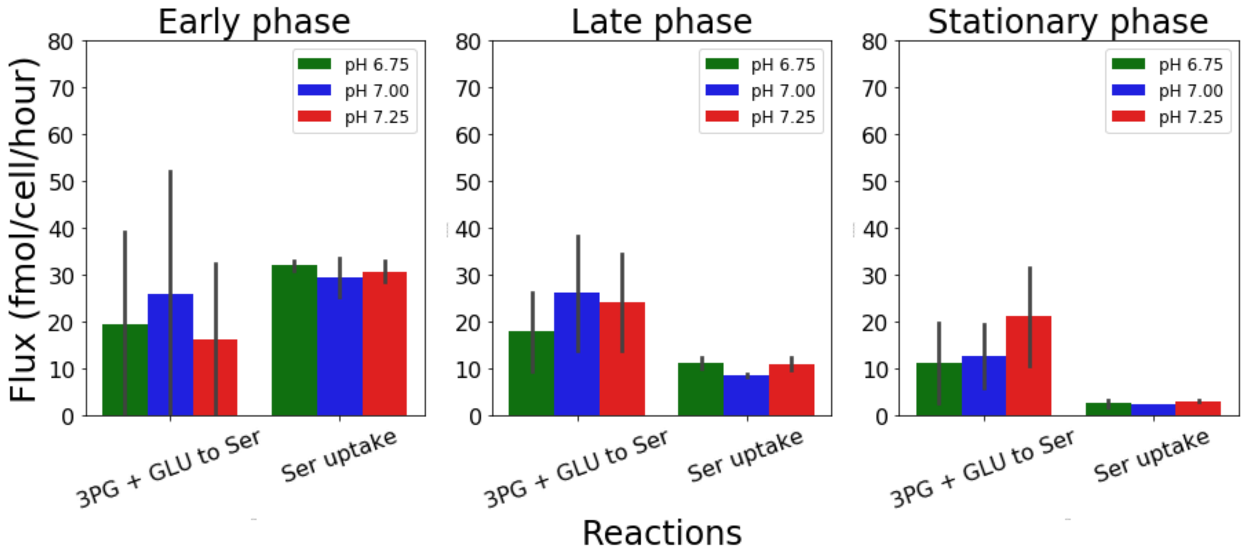


Figure S3: FVA identifies the sources of intracellular serine production during different phases of the culture at the three different pH conditions.

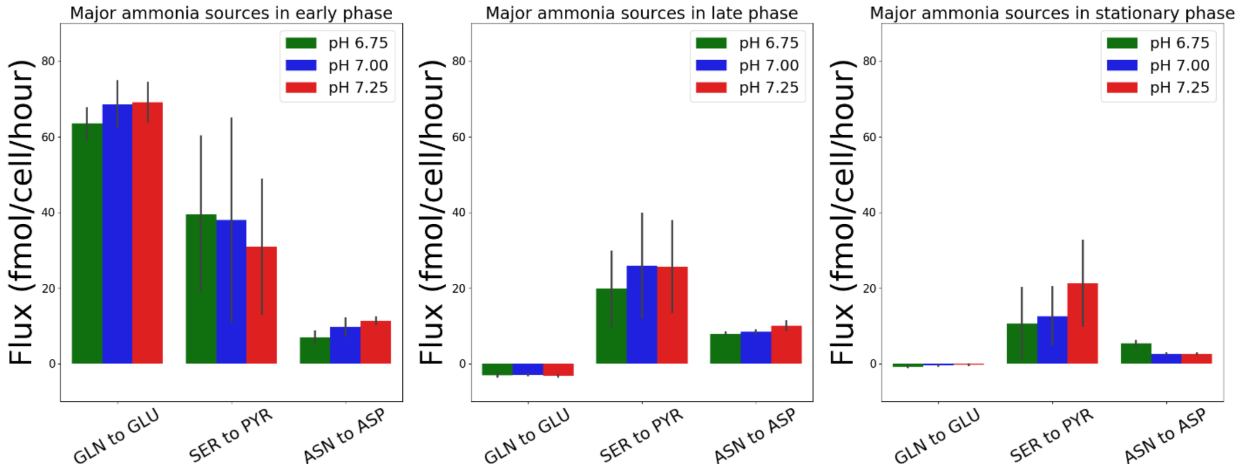


Fig. S4: Major sources of intracellular ammonia production during different culture phases at the three different pH conditions.

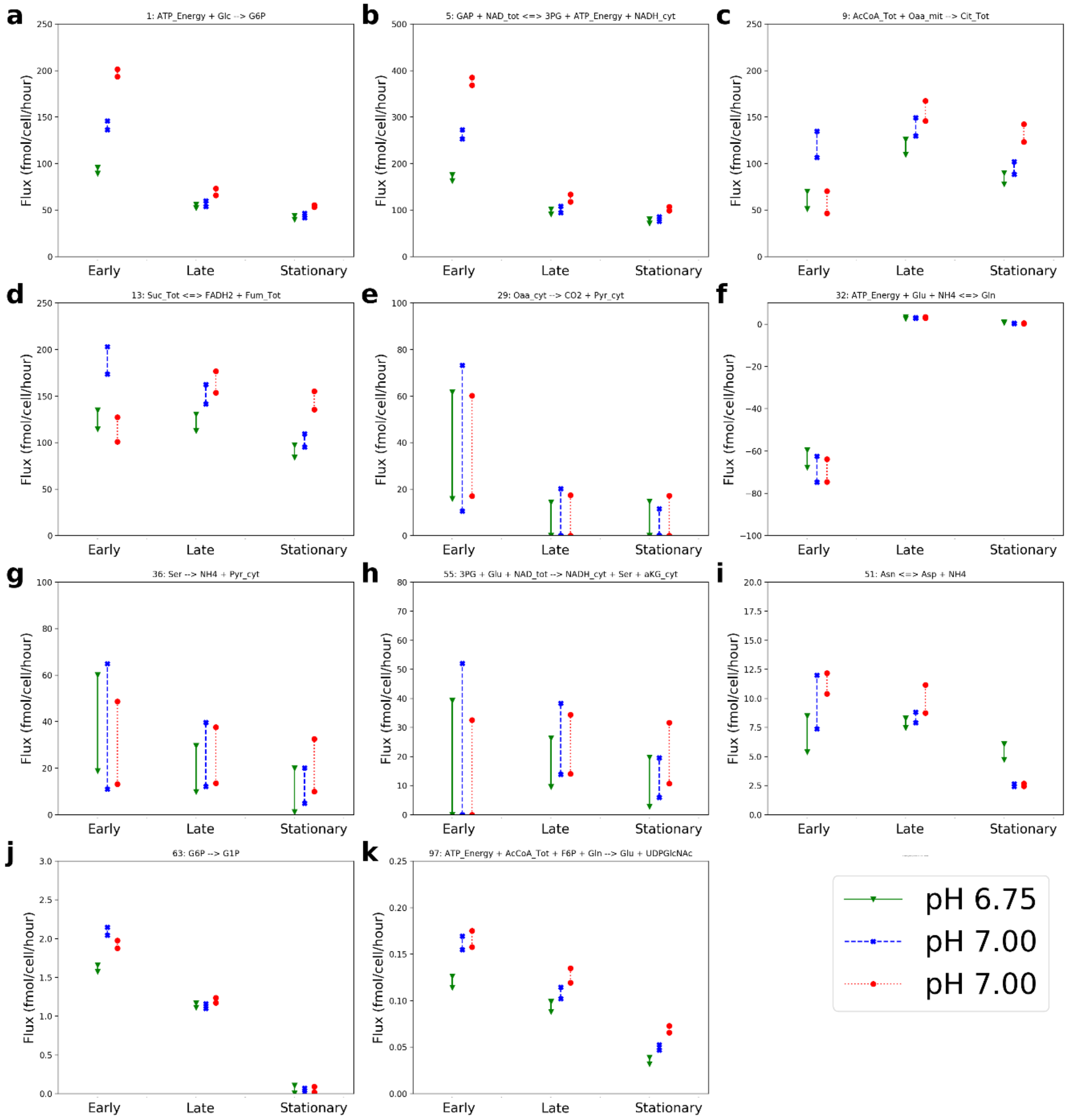


Fig. S5: Flux variability analysis (FVA) shows that bioreactor pH and culture duration can impact glycolysis, oxaloacetate to pyruvate fluxes, conversion of serine to pyruvate, biomass precursors, nucleotide sugar synthesis rates and amino acid metabolism (asparagine, aspartate, glutamate, and glutamine).

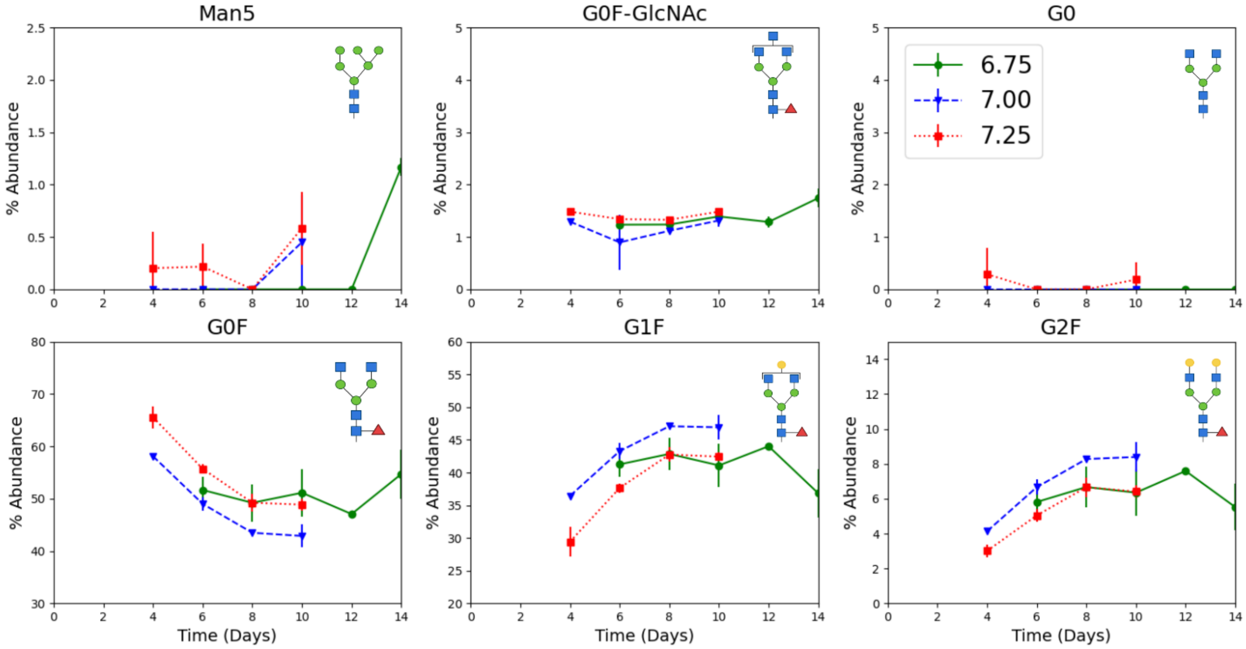


Fig. S6: Dynamic N-linked glycosylation data for Fc region for all the detected glycans.

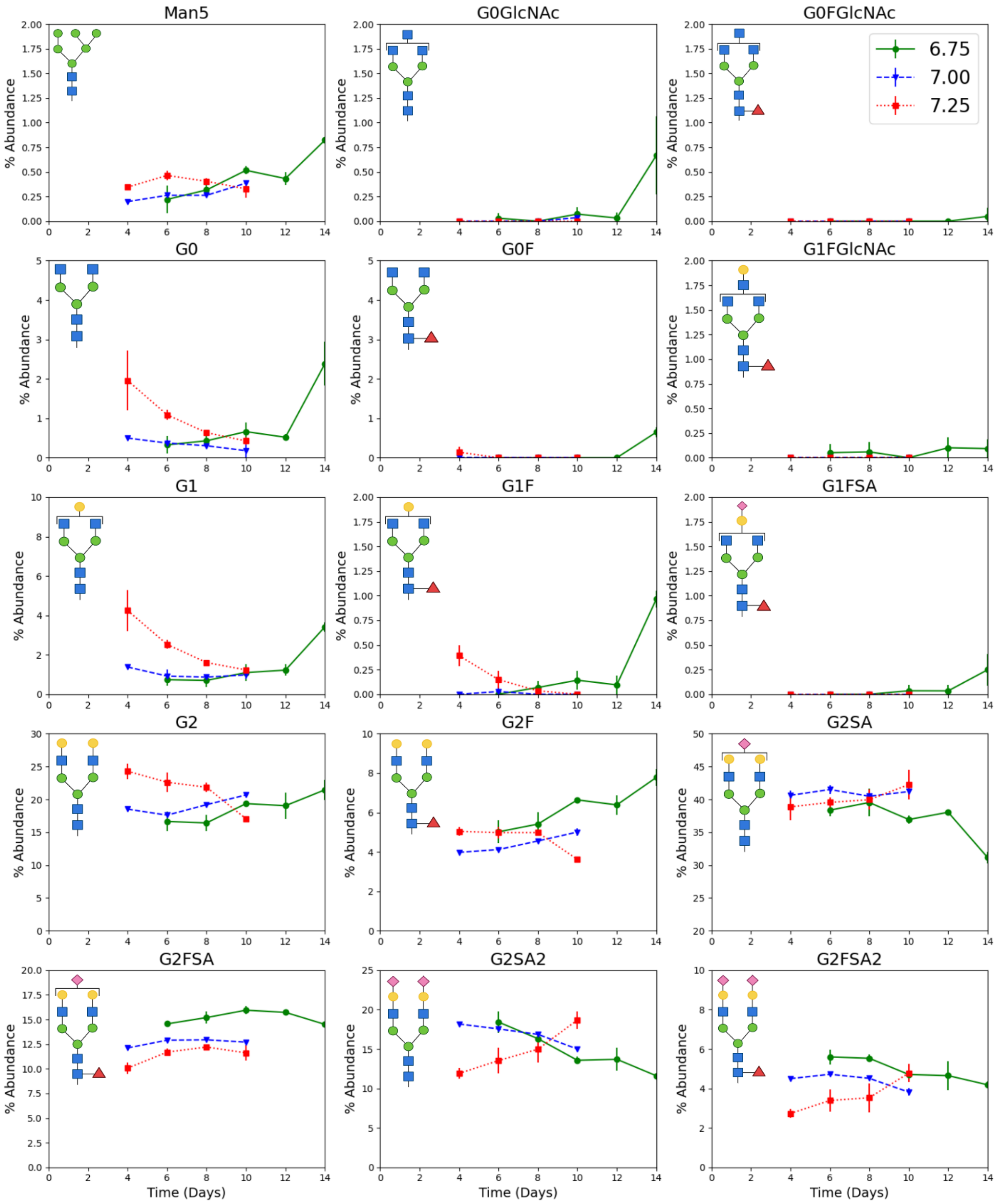


Fig. S7: Dynamic N-linked glycosylation profiles for all the glycans detected in the Fab region.
